# Nutritional aspects of honey bee-collected pollen and constraints on colony development in the eastern Mediterranean

**DOI:** 10.1101/008524

**Authors:** Dorit Avni, Harmen P. Hendriksma, Arnon Dag, Zehava Uni, Sharoni Shafir

## Abstract

**\*Graphical Abstract (for review):** 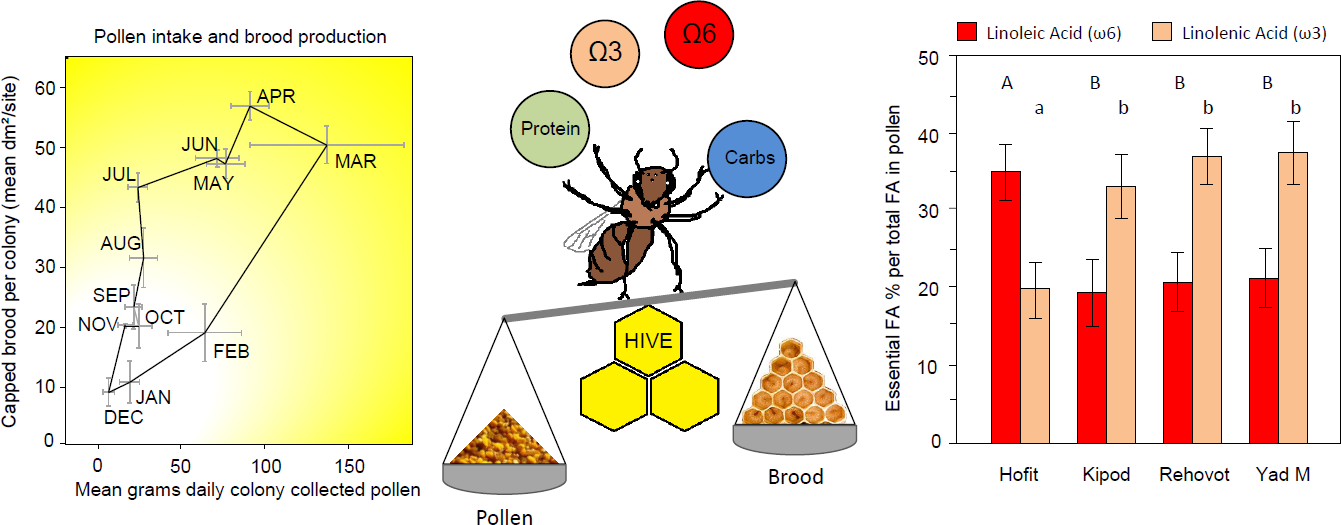

**\*Highlights (for review):**

- Honey bee colonies in Israel collect a mean 7 kg protein and 0.7 kg fat per year
- At maximal colony development, honey bee queens lay up to 3,300 eggs per day
- Amount and content of collected pollen differ between sites and seasons
- Pollen linoleic acid and protein levels best describe bee production cost
- Colonies seem limited first by climate, then protein and finally specific nutrients

**ABSTRACT:** Pollen is the main protein and lipid source for honey bees (*Apis mellifera*), and nutritionally impoverished landscapes pose a threat to colony development. To determine colony nutritional demands, we analyzed a yearly cycle of bee-collected pollen from colonies in the field and compared it to colony worker production and honey bee body composition, for the first time in social insects. We monitored monthly bee production in ten colonies at each of seven sites throughout Israel, and trapped pollen bi-monthly in five additional colonies at each of four of these sites. Pollen mixtures from each sampling date and site were analyzed for weight, total protein, total fatty acids (FAs), and FA composition. Compared to more temperate climates, the eastern Mediterranean allows a relatively high yearly colony growth of ca. 300,000 to 400,000 bees. Colonies at higher elevation above sea level showed lower growth rates. Queen egg-laying rate did not seem to limit growth, as peaks in capped brood areas showed that queens lay a prolific 2,000 eggs a day on average, with up to 3,300 eggs in individual cases. Pollen uptake varied significantly among sites and seasons, with an overall annual mean total 16.8 kg per colony, containing 7.14 kg protein and 677 g fat. Overall mean pollen protein content was high (39.8%), and mean total FA content was 3.8%. Production cost, as expressed by the amount of nutrient used per bee, was least variable for linoleic acid and protein, suggesting these as the best descriptive variables for total number of bees produced. Linolenic acid levels in pollen during the autumn were relatively low, and supplementing colonies with this essential FA may mitigate potential nutritional deficiency. The essentiality of linoleic and linolenic acids was consistent with these FAs’ tendency to be present at higher levels in collected pollen than in the expected nutrients in bee bodies, demonstrating a well-developed adjustment between pollinator nutritional demands and the nutritional value of food offered by pollinated plants.

## 1. Introduction

Honey bees collect pollen from a wide range of flowering plants, which usually fulfills their dietary requirements for proteins, lipids, minerals, and vitamins. When pollen is abundant, a honey bee colony collects between 15 and 55 kg pollen annually (Winston, 1987). Protein levels in the range of 2.5% to 61% have been reported in honey bee-collected pollen (de Arruda et al., 2013; Odoux et al., 2012; Roulston and Cane, 2000; Schmidt, 1984; Schmidt and Buchmann, 1999; Yang et al., 2013). Protein is essential for proper development and function of body tissue, muscles, membranes and glands (Herbert, 1999). Variability in pollen quantity and quality may pose a challenge for nutritional homeostasis within the colony. A nutritional deficit may shorten life span, reduce brood production, inhibit gland development, enhance emergence of diseases and reduce honey bee weight (Alaux et al., 2010; Schmidt, 1984; Schmidt et al., 1987; Schmidt et al., 1995).

Honey bee foragers collect nectar from plants to produce honey, which is the main source of carbohydrate nutrition for the colony. Honey is the alimentary resource for adult honey bees and the developing brood. In addition, the carbohydrates are used as fuel to thermoregulate the brood nest and as flight fuel for foragers. Carbohydrates are also used by young honey bees to biosynthesize lipids. For example, honey bees produce wax, which is the main building material for honeycombs. In Israel, honey bee hives yield 24.3 to 31.3 kg honey annually (Avni et al., 2009). In more temperate regions, honey bee colonies can collect up to 200 kg honey in a year (Winston, 1987).

Pollen is an essential source of lipids for honey bees, used for energy and as a structural component of cell membranes. Fatty acid (FA) content of pollen ranges between 1% and 20%, depending on the plant species (Brodschneider and Crailsheim, 2010). Most insects require essential polyunsaturated fatty acids (PUFA) in their diet, and linoleic and linolenic acids usually satisfy this nutritional need (Canavoso et al., 2001; Cohen, 2004; Khani et al., 2007; Nurullahoglu et al., 2004; Wang et al., 2006). These FA have multiple double bonds, starting at the 3^rd^ and 6^th^ carbon atom from the omega group end, hence alternatively named omega 3 (18:3ω3 / C18:3 cis-9,12,15 / linolenic acid) and omega 6 (18:2ω6 / C18:2 cis-9,12 / linoleic acid).

Pollen is typically rich in water-soluble vitamins, and it provides bees with their vitamin requirements. In general, as long as pollen is abundant, the honey bee requirements for these nutrients are satisfied (Brodschneider and Crailsheim, 2010; Herbert, 1999). Honey bee mineral requirements are the least known of all nutritional components (Cohen, 2004). Minerals are also obtained by the bees from pollen, which contains ash at 2.5% to 6.5% of its dry weight. Addition of 1% pollen ash to a synthetic honey bee diet increased brood rearing significantly, but exceeding the ash content by more than 2% did not seem to be advantageous (Herbert and Shimanuki, 1978).

Nutritional ecology is a rapidly growing field, especially for bee species; a nutritional deficiency due to anthropogenically altered landscapes could have implications for the global decline in bee populations. Suboptimal nutrient balance has been argued to be one of the threats to pollinator populations (Vanbergen et al., 2013). The division of labor in eusocial insect colonies adds a level of nutritional complexity. In the honey bee colony, older foragers collect the food, which is consumed by younger workers who then feed the larvae. Thus, the body composition of emerging adults is indirectly affected by the diet collected by the colony.

In the current study, analyses of pollen content and honey bee body composition were used to determine honey bee colony nutritional demands. This method is commonly used by animal nutritionists to determine the animal’s nutritional demands and compose a suitable artificial diet to satisfy them (Kratzer et al., 1994; Leeson and Summers, 2001; Rock and King, 1967). Here, we applied this method for the first time to a eusocial insect.

Long-term studies of honey bee colony pollen collection and nutrition can serve to elucidate the timing of major colony life-cycle events, such as growth, reproductive swarming, migration, and senescence. The current objective was to quantify annual amounts and composition of honey bee-collected pollen (total protein, total FA, and FA profiles), and compare them to the colony’s performance in terms of population growth and honey bee body composition. Our first hypothesis was that temporal and spatial effects govern the nutritional contents and amounts of honey bee-collected pollen. Secondly, we expected honey bee colony growth to be affected by altitude (as a measure of the growth season), and by pollen quantity and nutritional quality.

## 2. Materials and methods

### 2.1. Sites

Honey bee colonies (*Apis mellifera ligustica*) were monitored between November 2004 and December 2005 at geographically distinct sites (Fig. 1). These locations had different habitats and surrounding flora representing the Mediterranean climate (Avni et al., 2009), which is subtropical with long, hot and dry summers and relatively short, cool, rainy winters (Köppen climate classification Csa). The sites differed in elevation above sea level and rainfall (Table 1).

**Fig. 1.**
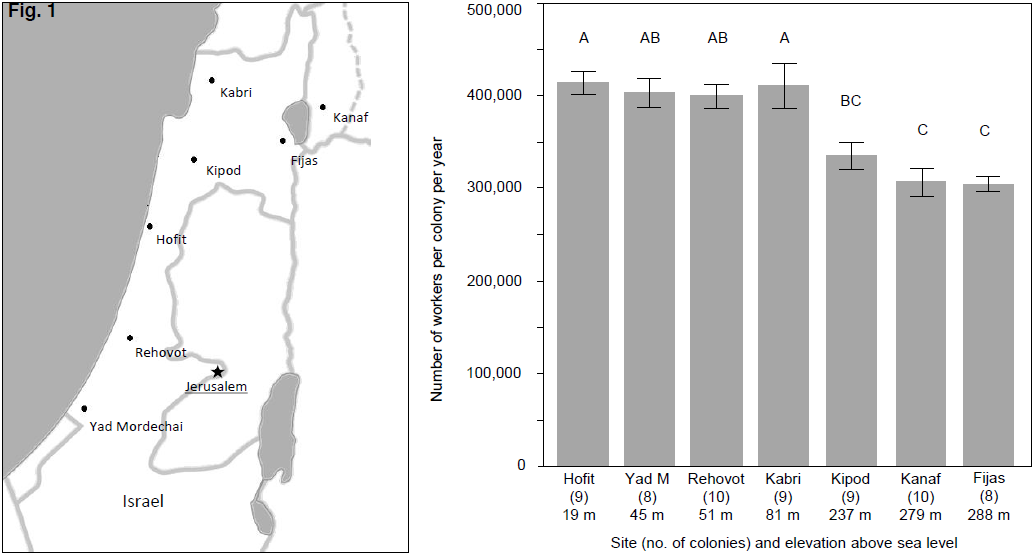
Honey bee colony development was monitored throughout Israel at seven sites, indicated on the map with closed circles (left pane). Mean (± SE) number of bees born in a honey bee colony over one year, as based on capped brood cell dynamics in 63 colonies (right pane). Sites are ordered according to increasing altitude, which correlates negatively with population growth per colony (linear regression: *R*^2^ = 0.50, *F*_1,61_ = 42.64, *p* < 0.001).

**Table 1.** The experimental sites where honey bee colonies were monitored. Rainfall data were derived from the nearest weather stations (data.gov.il/ims), with reported total rainfall for the season 2004–2005.

| Site | Hofit | Yad M. | Rehovot | Kabri | Kipod | Kanaf | Fijas |
| --- | --- | --- | --- | --- | --- | --- | --- |
| Latitude N | 32°23'37" | 31°35'25" | 31°54'26" | 32°01'15" | 32°36'18" | 32°52'11" | 32°41'43" |
| Longitude E | 34°52'30" | 34°34'07" | 34°48'16" | 35°08'50" | 35°07'20" | 35°41'52" | 35°33'05" |
| Rain (mm) | 556 | 433 | 466 | 597 | 1162 | 696 | 902 |
| Elevation (m) | 19 | 45 | 51 | 89 | 237 | 279 | 288 |

### 2.2. Honey bee colonies and assessments

Fifteen honey bee colonies were kept at each of the seven sites. At the beginning of the experiment, test colonies had on average 6.5 populated frames and 25 dm^2^ brood (Supplement 1). Young queens were introduced in the autumn, and all were daughters of the same mother-queen from the local breeding program of the Ministry of Agriculture apiary in Tzrifin, Israel. The queens were naturally mated with local drones from Italian strains. Local beekeeping practice was followed, applying swarm-suppressing measures such as the use of young queens, the removal of developing queen cells, and the placement of additional hive boxes on top of the brood frames to increase the amount of space available for food storage and brood production.

Every month, we assessed 10 hives per site for comb surface areas of brood and pollen and honey storage (Supplement 1). The areas were estimated by dividing each side of each comb into eight squares (10 cm × 10 cm = 1 dm^2^), and counting the number of squares of each type (Kalev et al., 2002). The growth of each colony over one year was determined by taking the sum of the daily capped brood amount (see section 2.3), multiplying it by the number of cells per dm^2^ (397 worker cells; based on counting the total cell number on a comb foundation), and dividing by 12, the number of days that brood cells are capped (Winston, 1987). From a total 70 colonies (*n*_observations_ = 802), seven colonies lost their queen during the year and were excluded from population growth analyses. The peak in capped brood area per colony (on any one of the sample days) was used to indicate the queen’s egg-laying capacity. For example, a capped brood area of 60.5 dm^2^ contains 24,000 worker cells (397 cells·dm^-2^), thus considering the 12 days during which the brood is capped, the queen must have ovipositioned an average 2,000 eggs per day in the preceding period.

Bottom pollen traps were placed under five hives at each site. As pollen trapping may influence colony development, the traps were not placed under the hives used for colony growth monitoring. The traps consisted of a double net with 0.5 cm × 0.5 cm openings that allowed the bees to pass through, while trapping the pollen pellets on their hind legs. Below the nets there was a drawer to collect the fallen pollen pellets. The percentage of incoming pellets captured by the traps was checked in a separate experiment: mean (± SE) capture rate was 40.3% ± 3.2, *n* = 33 trials with six colonies (Avni et al., 2009). The traps were closed for 3 to 5 days every 2 weeks, and the captured pollen was collected, weighed to the nearest 0.01 g, and stored at −20 °C until analysis.

At four of the seven sites (Hofit, Yad Mordechay, Rehovot, Kipod), on each sampling date, a spoon of trapped pollen from each of the five colonies was pooled into a single sample for nutritional analysis. Each sample was mixed and crushed mechanically by mortar and pestle (Pirk et al., 2009), and 2–3 g were used for protein and FA analyses (see section 2.5).

Colonies’ daily pollen uptake was calculated as follows: the collected pollen weight per sample was divided by the number of days of trapping activity (3–5), and divided by 40.3% to correct for the previously reported trap efficiency. Overall, a mean 15.3 samples were taken per colony, with the pollen traps active for a total of 52 days ± 2 SE per colony, over a total mean period of 305 days ± 2 SE (*n*_colonies_ = 20, *n*_sites_ = 4).

### 2.3. Annual interpolation

The sample dates for pollen and brood differed within and between sites. To enable comparisons over time, and to estimate yearly totals, the data were interpolated between sampling dates. For each of the test colonies, we fit a kernel smooth curve at α level of 0.3 for the relationship between amount of capped brood (*n* = 63 colonies) or amount of bee-collected pollen (*n* = 20 colonies) and date, yielding a coefficient of determination (*R*^2^) for each colony. The means of these coefficients were *R*^2^_brood_ = 0.86 and *R*^2^_pollen_ = 0.72, respectively. To cover a full-year cycle, the first brood data points were repeated after 365 days, and the pollen data were set to zero on 1 Dec 2004 and 2005. In addition, the mean amounts of collected pollen (five colonies per sampling date) were multiplied by data on sample content (see section 2.5), and thereafter interpolated to gain estimates on annual colony intake (mean *R*^2^ = 0.70).

### 2.4. Honey bee body composition

To assess body composition of bees from nutritionally nondeprived colonies, we kept six colonies in a netted enclosure and fed them pollen patties (50% multifloral pollen mixed with 50% sugar, w/w) ad libitum. Emerging bees were marked and returned to the colonies to be collected 8 days later. The body weight (*n* = 212 bees at 8 days of age), protein content and FA composition were measured (*n* = 12; 6 colonies × 2 bees) and used as a reference for expected body composition of field bees (see section 2.5). A worker honey bee is often considered an adult after emergence from the cell, though during the first 6 days, substantial growth occurs, expressed as an increase in body weight and a 25-50% increase of total protein content of (Haydak, 1934). We therefore analyzed the body composition of 8-day-old adults.

### 2.5. Protein and FA analyses

Pollen and honey bee samples were homogenized with water (0.1 g in 1.0 ml water) in a Mini Bead Beater (Biospec Products, Bartlesville, OK, USA). Samples from the homogenate (0.1 ml) were diluted 1:5 with water and total protein (mg/g) was detected by Bio-Rad protein assay (Bradford, 1976), which uses a copper-based reagent to stain protein. To quantify the proteins, bovine serum albumin was used as a standard.

For total FA analysis, 0.1 ml of homogenate (0.1 g in 1 ml water) was subjected to basic hydrolysis, followed by petrol ether extraction. Nonsaponified matter was removed. The sample was then transferred to an acidic environment for FA release. A second petrol ether extraction was conducted, followed by evaporation and methylation with 5% sulfuric acid in methanol. Then a third petrol ether extraction was conducted for methyl ester collection. The methyl esters were separated by HP 5890 gas chromatograph with FID. An internal standard (heptanoic acid, C17:0, Sigma, Israel) was added to each sample to quantify FA amounts (Sklan and Budowski, 1979). The injector, oven and detector temperatures were 230 °C, and constant 180 °C and 235 °C, respectively. We used a 1/8”, 2.5 m stainless-steel packed column filled with GP 10% SP-2330 on 100/120 chromosorb WAW (Supelco).

### 2.6. Data analysis

Values of the monitored colony parameters (honey bee colony growth, pollen collection, relative levels of pollen constituents) were compared by ANOVA with Tukey’s honestly significant difference test (JMP, Version 10, SAS Institute Inc.). Pollen data were analyzed on two explanatory variables, season (4 levels) and site (4 levels), by two-way ANOVA. The main results were considered statistically significant at *p*-values below α = 0.05. The pollen fat-content variables (total FA, linolenic and linoleic acids) were not independent and thus tested at α = 0.01. The cost of honey bee production was calculated by dividing the total amount of nutrients taken up by the number of honey bees produced. We expected nutrients that govern or limit production to show a stable cost per bee; such nutrients were identified by a low coefficient of variation (CV) for the honey bee production cost.

## 3. Results

The number of new bees produced per colony during the year differed among the seven sites, with a mean range of ca. 300,000 to 400,000 (Fig. 1; *F*_6,56_ = 10.2, *p* < 0.001), a minimum of 217,000 and a maximum 520,000 bees per year (*n* = 63 colonies). Total honey bee production was lower at higher elevations (Fig. 1; *R*^2^ = 0.50, *F*_1,61_ = 42.64, *p* < 0.001). The overall egg-laying rate of queens approximated 1,000 eggs per day (on average 367,000 bees ± 8,000 SE in a year). The mean of yearly peak for capped brood area in the 63 colonies was 62.2 dm^2^, corresponding to the queens laying approximately 2,078 eggs per day during peak periods. The two colonies with the highest peaks had capped broods of 99.5 dm^2^ and 89.5 dm^2^, with queens laying 3,300 and 3,000 eggs per day, respectively.

The amount of pollen collected by the honey bees in Israel varied over the seasons (Table 2). In the absence of rain, pollen collection decreased over the summer, with the smallest amounts collected in the autumn (Fig. 2). Pollen collection also differed among sites (Table 2), with an annual colony mean of 16.8 kg. The smallest amount of pollen was collected in Hofit (insignificant by post-hoc test), compared to Yad Mordechai, Rehovot, and Kipod (Table 2). Similarly significant was the site effect for the yearly amount of collected pollen (*F*_3,16_ = 3.48, *p* = 0.04), albeit with a post-hoc difference between the lowest and highest levels (Hofit and Yad Mordechai, respectively, Table 3).

**Fig. 2.**
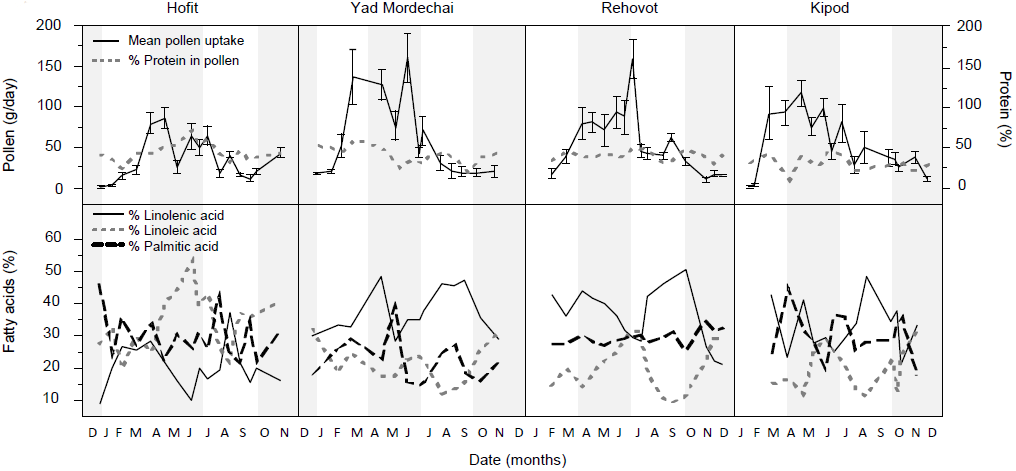
Amount of bee-collected pollen and its relative protein content over time, at four sites in four seasons (per solstice and equinox, as indicated by the white and gray backgrounds). In the upper row, left axis shows weight of collected pollen; means with SE (error bars) are from five colonies; right axis shows protein percentage in pollen samples. The lower row shows the three most abundant fatty acids, as percentage of the total FA amount in pollen. Note the reduction in Hofit of both amount of pollen collected over the year, and contrasting levels of linolenic and linoleic acids, in comparison to the other sites.

**Table 2.**
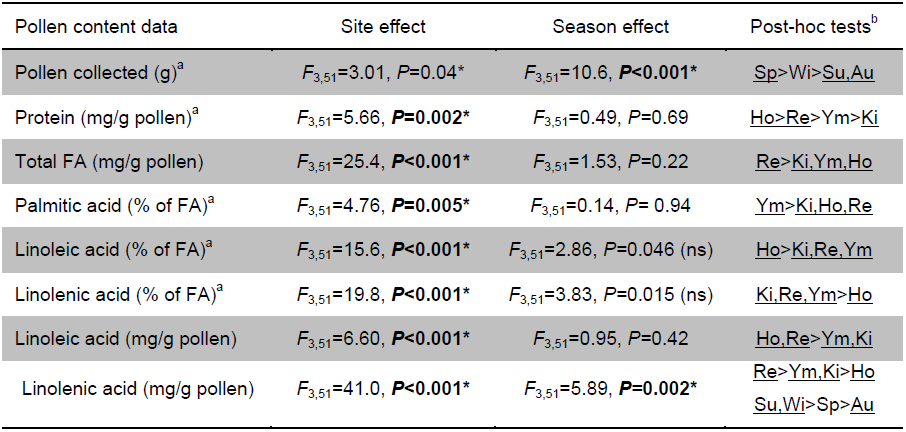
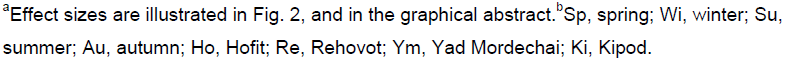
Two-way ANOVA of bee-collected pollen quantity and contents over time and place. *Statistically significant considering a threshold at α = 0.05, or for the interdependent fatty acid (FA) tests at α = 0.01. For all significant results in bold, post-hoc results are indicated (right column). Differences exist between the underlined groupings. The indicated site effect on collected pollen was not post-hoc significant. Note: Pollen amount is strongly influenced by season, and pollen contents is typically influenced by site.

| Pollen content data | Site effect | Season effect | Post-hoc tests<sup>b</sup> |
| --- | --- | --- | --- |
| Pollen collected (g)<sup>a</sup> | $F_{3,51}=3.01$, $P=0.04^*$ | $F_{3,51}=10.6$, $P<0.001^*$ | <u>Sp>Wi>Su,Au</u> |
| Protein (mg/g pollen)<sup>a</sup> | $F_{3,51}=5.66$, $P=0.002^*$ | $F_{3,51}=0.49$, $P=0.69$ | <u>Ho>Re>Ym>Ki</u> |
| Total FA (mg/g pollen) | $F_{3,51}=25.4$, $P<0.001^*$ | $F_{3,51}=1.53$, $P=0.22$ | <u>Re>Ki,Ym,Ho</u> |
| Palmitic acid (% of FA)<sup>a</sup> | $F_{3,51}=4.76$, $P=0.005^*$ | $F_{3,51}=0.14$, $P=0.94$ | <u>Ym>Ki,Ho,Re</u> |
| Linoleic acid (% of FA)<sup>a</sup> | $F_{3,51}=15.6$, $P<0.001^*$ | $F_{3,51}=2.86$, $P=0.046$ (ns) | <u>Ho>Ki,Re,Ym</u> |
| Linolenic acid (% of FA)<sup>a</sup> | $F_{3,51}=19.8$, $P<0.001^*$ | $F_{3,51}=3.83$, $P=0.015$ (ns) | <u>Ki,Re,Ym>Ho</u> |
| Linoleic acid (mg/g pollen) | $F_{3,51}=6.60$, $P<0.001^*$ | $F_{3,51}=0.95$, $P=0.42$ | <u>Ho,Re>Ym,Ki</u> |
| Linolenic acid (mg/g pollen) | $F_{3,51}=41.0$, $P<0.001^*$ | $F_{3,51}=5.89$, $P=0.002^*$ | <u>Re>Ym,Ki>Ho</u><br><u>Su,Wi>Sp>Au</u> |

<sup>a</sup>Effect sizes are illustrated in Fig. 2, and in the graphical abstract.<sup>b</sup>Sp, spring; Wi, winter; Su, summer; Au, autumn; Ho, Hofit; Re, Rehovot; Ym, Yad Mordechai; Ki, Kipod.

**Table 3.** Estimates per year in mean ± SE (no. of colonies in parenthesis), or total estimate per site. The overall number of bees produced is given for the seven sites. For four sites, the average yearly total pollen amount and the totals for protein and FA contents are listed. Pollen content data originate from pooled batches of five sampled colonies per site, collected every month. The absolute uptake of each nutrient was interpolated over a period of 1 year to assess the total uptake of nutrients (protein and FAs).

| Estimate | Overall mean | Hofit | Yad Mordechai | Rehovot | Kipod |
| --- | --- | --- | --- | --- | --- |
| Bees * 1,000 | 367 ± 8.1 (63) | 413 ± 14 (9) | 402 ± 15 (9) | 399 ± 13 (10) | 335 ± 14 (9) |
| Pollen (kg) | 16.8 ± 1.2 (20) | 11.8 ± 0.7 (5) | 20.9 ± 2.9 (5) | 16.9 ± 2.2 (5) | 17.6 ± 1.6 (5) |
| Protein (kg) | 7.14 | 5.88 | 9.50 | 7.06 | 6.13 |
| Total FA (g) | 677 | 358 | 689 | 926 | 735 |
| Palmitic acid (g) | 190 | 104 | 164 | 269 | 225 |
| Linoleic acid (g) | 152 | 129 | 146 | 199 | 135 |
| Linolenic acid (g) | 234 | 74 | 243 | 352 | 244 |

In contrast, the content of pollen constituents was stable over the seasons, but differed significantly among sites (Table 2). In the 58 pollen samples, the protein content was on average 39.8%, ranging between a minimum 10.6% and maximum 73.0%. The protein levels differed significantly with a mean 30.4% in Kipod, 40.5% in Yad Mordechai, 40.9% in Rehovot and 45.8% in Hofit (Table 2, Fig. 2, and see also graphical abstract). Among sites, the lowest annual protein uptake was in Hofit (<6 kg), despite its highest annual bee production (Table 3).

Total FA content in the pollen averaged 3.8% (38.4 mg/g pollen), ranging from a minimum 2.3% to a maximum 6.6% (*n* = 58 samples). There was a difference between sites (Table 2), with means of 3.2%, 3.3% and 3.6% in Hofit, Yad Mordechai and Kipod, respectively, and a significantly higher content of 5.1% in Rehovot (see also graphical abstract). On average, a colony collected about ten times more protein than fat per year (Table 3).

Per total amount of FA in the pollen, the three major FAs were palmitic acid (16:0), linoleic acid (18:2ω6) and linolenic acid (18:3ω3), with mean (± SE) values of 28.3% ± 6.9, 24.7% ± 9.7 and 31.6% ± 10.7, respectively (Fig. 2). Together, these three FAs made up over 80% of the total FAs. Other FAs detected were: myristic, palmitoleic, stearic, oleic and arachidonic acids (Table 4). The level of palmitic acid remained relatively stable throughout the year, whereas linoleic and linolenic acids showed more fluctuations (Fig. 2). The greater stability of palmitic acid levels (as % in all pollen samples) was expressed in their lower CV value of 24.4, in comparison to 39.1 and 33.8 for the essential FAs linoleic and linolenic acid, respectively. The variability in linoleic and linolenic acids was partly due to a striking difference between Hofit and the other three sites (Fig. 2). In Hofit, linolenic acid levels were typically low relative to linoleic acid, whereas the opposite pattern was found at the other sites (Graphic abstract, Table 4).

**Table 4.** Fatty acid (FA) content in worker bee bodies and in pollen samples at four different sites. Data are presented as the percentage of total FA content and are reported as means ± SE (number of samples analyzed is in parentheses). Yearly variation for palmitic, linoleic and linolenic acids is illustrated in Fig. 2, and in the graphical abstract. After the trivial names, the lipid numbers are given, indicating the number of carbon atoms and the number of double bonds in the fatty acid.

| Fatty acid |  | Bees (12) | Hofit (16) | Yad M. (13) | Rehovot (16) | Kipod (13) |
| --- | --- | --- | --- | --- | --- | --- |
| Myristic acid | 14:0 | 2.05 ± 0.11 | 0.83 ± 0.11 | 1.10 ± 0.11 | 1.19 ± 0.14 | 0.83 ± 0.11 |
| Palmitic acid | 16:0 | 18.57 ± 1.11 | 30.0 ± 1.79 | 22.3 ± 1.96 | 30.0 ± 0.58 | 30.1 ± 2.05 |
| Palmitoleic acid | 16:1 | 2.36 ± 0.24 | 0.07 ± 0.07 | 0.36 ± 0.10 | 0.42 ± 0.16 | 0.27 ± 0.07 |
| Stearic acid | 18:0 | 12.54 ± 0.61 | 3.59 ± 0.41 | 5.30 ± 0.52 | 2.97 ± 0.28 | 4.60 ± 0.33 |
| Oleic acid | 18:1 | 31.28 ± 0.90 | 9.84 ± 0.89 | 10.25 ± 0.82 | 5.67 ± 0.49 | 9.73 ± 0.58 |
| Linoleic acid | 18:2ω6 | 11.43 ± 0.72 | 34.6 ± 2.26 | 21.4 ± 1.73 | 21.4 ± 1.93 | 20.0 ± 1.90 |
| Linolenic acid | 18:3ω3 | 21.76 ± 1.06 | 20.3 ± 1.73 | 37.2 ± 2.00 | 37.4 ± 2.39 | 32.8 ± 2.22 |
| Arachidonic acid | 20:4 | 17.04 ± 0.99 | 0.77 ± 0.16 | 2.14 ± 0.31 | 0.98 ± 0.13 | 1.72 ± 0.22 |

The mean (± SE) honey bee weight, and protein and total FA contents of 12 bees reared under controlled conditions were 111.4 mg ± 3.2, 11.1% ± 0.32 and 1.7% ± 0.10, respectively. Fig. 3 shows the total protein and FA contents in the pollen collected by the colonies at each site during the year, and total expected amounts in honey bee bodies based on the number of bees produced during the year (panes A and B, respectively). At all sites, the amount of protein in the collected pollen was greater than that in the bees’ bodies. For FAs, however, at some sites the content in the bees was greater than that in the collected pollen.

**Fig. 3.**
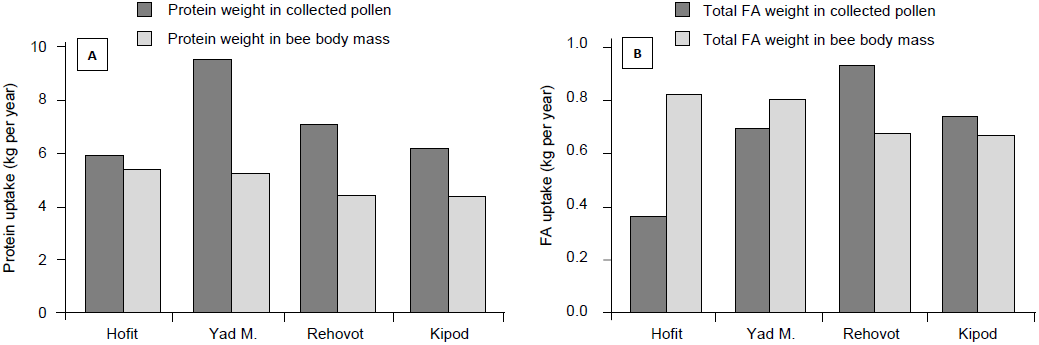
Comparison in four sites in Israel between protein (A) and total fatty acid (FA) (B) levels in pollen collected by colonies and those calculated to be present in the total produced bees’ body mass. Note that the honey bee protein is accounted for, with more total protein in pollen than in honey bee bodies, whereas FAs were also derived by means other than pollen content, for instance by biosynthesis.

Table 4 shows the individual FA contents of bee bodies and collected pollen. In addition, Fig. 4 compares the FA balance between bee bodies and collected pollen. In general, there was a positive FA budget for palmitic, linoleic and linolenic acids, with more FA in the collected pollen than in the bee bodies. Hofit was the exception, with negative budgets for palmitic (nonessential) and linolenic (essential) acids. Other nonessential FAs were more abundant in the bees than in the pollen that they had collected. From largest to smallest difference, these included oleic, stearic, palmitoleic, and myristic acids.

**Fig. 4.**
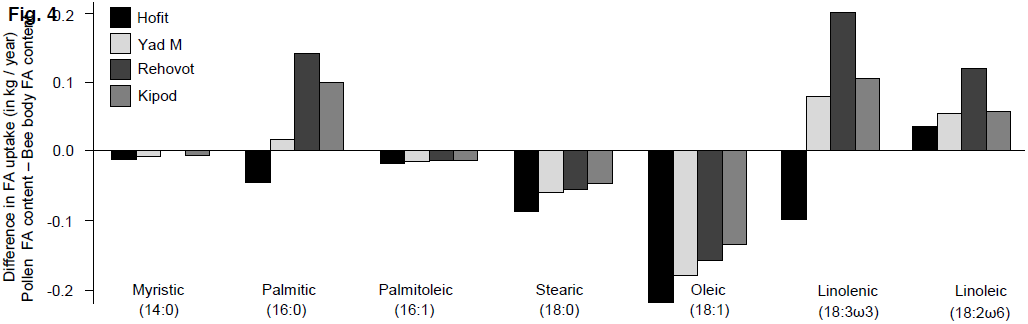
Fatty acid (FA) uptake by colonies over a whole year was compared with the expected FA composition of all bees produced in a year. The difference between colony FA intake and output was calculated by subtracting the honey bee body FA contents from the pollen FA contents. Thus, a positive value indicates that more pollen FA was taken in than was used for the production of bees. A negative value shows that pollen contained less of this FA compared to its output in the body mass, indicating its biosynthesis within the colony. Note that the essential FAs were available in sufficient quantities, with positive linolenic and linoleic acid values, except in Hofit. The lipid numbers are given in parentheses, indicating the number of carbon atoms and the number of double bonds in the fatty acid.

Totaled over the year, the brood production cost was on average 51.3 mg pollen per produced honey bee (Table 5). In absolute uptake of pollen contents, the cost showed the least variation for linoleic acid and protein (lowest CVs; Table 5), and these were accordingly the best descriptive variables for the total number of workers produced, as compared to, for example, the gross amount of pollen or linolenic acid (Table 5).

**Table 5.** Nutritional costs of producing a honey bee at four sites in Israel, and overall mean. The last column is the coefficient of variation (CV = SD/mean), used to compare the costs at the four sites. The nutrients are listed according to increasing CV value. The nutrient with the lowest CV may be a limiting factor for growth, as it best describes the production cost of bees.

| Cost (nutrient per bee) | Overall mean | Hofit | Yad M. | Rehovot | Kipod | CV (%) |
| --- | --- | --- | --- | --- | --- | --- |
| mg linoleic acid | 0.46 | 0.37 | 0.42 | 0.58 | 0.47 | 20.0 |
| mg protein | 21.6 | 16.7 | 27.6 | 20.7 | 21.4 | 21.0 |
| mg pollen | 51.3 | 33.4 | 60.8 | 49.6 | 61.3 | 25.5 |
| mg total FA | 2.07 | 1.01 | 2.00 | 2.72 | 2.56 | 37.1 |
| mg palmitic acid | 0.59 | 0.29 | 0.48 | 0.79 | 0.78 | 41.5 |
| mg linolenic acid | 0.70 | 0.21 | 0.71 | 1.03 | 0.85 | 50.4 |

## 4. Discussion

Pollen fulfills honey bees’ nutritional demands for protein and fat. Whereas the importance of protein in pollen is well established, that of fat has often been overlooked (Cohen, 2004). Pollen protein and fat contents differ among plants. Here we measured the amount and content constituents of honey bee-collected pollens in Israel, and assessed the nutritional balance between the collected pollen and the number of bees produced in a colony. Amounts of collected pollen and brood both peaked in spring (see also graphical abstract). This fits the consensus that colony development and reproduction are mainly related to the amount of pollen collected, rather than its composition (Avni et al., 2009; Dimou and Thrasyvoulou, 2007; Schmidt and Buchmann, 1999). Nonetheless, we show in the present study that data on pollen content can be an excellent predictor of honey bee production levels, as compared to gross pollen amounts. The smaller CVs for pollen linoleic acid and protein contents suggested that these parameters best describe the overall amount of bees produced by colonies on a yearly basis (Table 5).

### 4.1. Total pollen collection and honey bee production

There was high seasonal variability in the amount of pollen collected by colonies (Table 2). Overall, colonies collected an annual average of 16.8 kg pollen. Similarly, colonies in Europe and North America have been estimated to collect between 13.4 kg and 55 kg pollen on a yearly basis (Crailsheim et al., 1992; Herbert, 1999; Odoux et al., 2012; Schmidt and Buchmann, 1999; Winston, 1987). Surprisingly, the average yearly production of 367,000 bees is notably high; double that of 117,000 to 150,000 worker brood cells per year (as listed for multiple studies in Table 5 within Crailsheim et al. (1992)), or 226,596 born worker bees (Page and Metcalf, 1984). A likely driver for the high annual brood-production level might lie in the subtropical Mediterranean climate in Israel, which allows for a prolonged brood-rearing season (Avni et al., 2009). High production, however, may be offset by shorter worker lifespan than in colonies in temperate climates, where winter (diutinus) workers are less active and live for several months (Amdam and Page, 2005).

At the height of colony growth in our study, the peak brood amounts indicated that queens laid, on average, 2,078 eggs per day, which is similar to the 1,950 to 2,500 eggs reported by Harbo (1986) based on brood area. Two colonies peaked with queens having laid 3,300 and 3,000 eggs per day, respectively, equivalent to 2 eggs per minute. Beekeepers have reported such observations anecdotally (Wright, 2008), but we believe that our data provide unprecedented support for the potentially high egg-laying rate of honey bee queens. This finding suggests that the queen’s maximum egg-laying capacity is not the limiting factor in yearly colony production, but that colony development is limited by other factors.

By dividing the colony’s annual pollen uptake by the number of bees produced, we calculated a pollen cost per produced honey bee (Table 5). The average cost of 51.3 mg pollen per honey bee produced in Israel reflects high efficiency compared to the previously reported range of 86 to 188 mg pollen per honey bee (Alfonsus, 1933; Babendreier et al., 2004; Hrassnigg and Crailsheim, 2005; Rosov, 1944; Wille et al., 1985). This efficiency may be partly attributed to the pollen content.

### 4.2. Protein content

The total amount of pollen collected by colonies in this study was in the low range of amounts reported for colonies in temperate regions; nevertheless, about twice as many bees were produced per year. This may be due to the high protein content of the pollen in Israel (39.8%), almost twice that reported in other studies. To compare with other Mediterranean countries, Tüylü and Sorkun (2006) reported an average 19.0% protein for 14 bee-collected monofloral pollens from Turkey, and Odoux et al. (2012) report a range of 16 to 29% protein for mixed bee-collected pollens from the south of France. In absolute protein amounts, Crailsheim et al. (1992) reported a yearly colony uptake of between 2.8 and 3.7 kg of protein per colony, whereas the estimated uptake in the current study was 7.14 kg. These figures illustrate how high protein intake might have enabled production of double the amount of bees with the same amount of pollen. The relatively high colony production levels in Israel in relation to the high protein levels could indicate that in more temperate regions, protein is a limiting nutrient.

Whereas pollen protein quantifications may differ according to the analytical method used (Roulston et al., 2000; Vanderplanck et al., 2014), between March and September 2005, 11 pollen samples from Amman, Jordan, were collected by Nizar Haddad’s group and analyzed by the same method presented in the current paper (Shafir et al., 2009): The average protein content was 28.7%, strengthening the finding of comparably higher pollen protein contents in Israel. Furthermore, since bees may add variable amounts of nectar to pollen pellets, protein percentage of honey bee-collected pollen may underestimate the protein percentage in pollen (Roulston et al., 2000). In the review by Roulston et al. (2000), hand-collected pollen from 62 bee-pollinated plants had an average 38.1% protein, as analyzed by the Bradford method, similar to our average of 39.8% protein found for 58 honey bee-collected pollen batches.

Importantly, the protein analyses in our study were internally consistent, rendering the comparative analyses, i.e. between seasons and sites, reliable. We found pollen protein content to be stable over the seasons, despite significant differences among sites (Table 2). The low pollen protein level in Kipod (see graphical abstract) may have contributed to the relatively low brood production at that site, in addition to the altitude effect which significantly reduced colony growth (Fig. 1).

### 4.3. FA content

We found a narrow range of 2.3% to 6.6% total FA in bee-collected pollen mixtures. Similarly, seven mixed bee-collected pollen samples of subtropical origin were reported to have a narrow range of 4.6% to 6.1% total FA (de Arruda et al., 2013), though mixed bee pollen of the western Mediterranean was reported to have a much higher and wider range of 7 to 24% lipids (Odoux et al., 2012).

Honey bees collect pollen of mixed sources, from an average six plants at any one time in Israel (Avni et al., 2009), and an average 11 plants in a subtropical region in Brazil (Hilgert-Moreira et al., 2014). A study by Schmidt et al. (1987) found that honey bee survival is optimal when feeding on a blend of five different pollens, compared to monofloral pollen diets. Hence, balancing by means of a polyfloral diet can be an active nutritional strategy for honey bees, to mitigate the risk of a particular nutritional source (a monofloral pollen diet) having a shortfall in essential nutrients. In addition, certain FAs, such as linoleic, linolenic, myristic and lauric acids, have bactericidal and antifungal properties that support colony hygiene (Manning, 2001; Manning and Harvey, 2002), and may therefore be important for disease resistance and colony survival.

We used the FA composition of bees fed a rich mixed pollen diet under controlled-enclosure conditions as a reference to assess honey bee nutritional demands. Although linolenic acid content in Hofit suggested a deficiency (Fig. 4), this site was not outperformed by the others in honey bee production (Fig. 1). It is possible that under deficient conditions, bees are produced that require fewer resources, for example, smaller bees produced by early capping (Schmickl and Crailsheim, 2001). It is also possible that microbial interactions, either in the gut or during pollen storage as bee bread, change the content of the honey bee’s food to compensate for the observed potential deficiencies (Douglas, 2013; Haydak and Vivino, 1950). However, in the absence of actual analyzed honey bee samples from each site, we have no means of explaining the discrepancy in linolenic acid levels in Hofit.

The essentiality of linoleic and linolenic acids is consistent with the fact that these FAs were generally (except for linolenic acid in Hofit) present at higher levels in the collected pollen than in the bees’ body mass (Fig. 4). The dominance of palmitic acid, a substrate for longer FA synthesis, and the two essential linolenic and linoleic acids in bee-collected pollen, demonstrates a well-developed adjustment between pollinator nutritional demands and the nutritional value of the food offered by pollinated plants.

### 4.4. Nutritional balancing

Some insects can balance their diet to compensate for a lack of essential nutrients (Dussutour and Simpson, 2009; Mayntz et al., 2005; Raubenheimer and Simpson, 1999; Simpson et al., 2004). We analyzed the nutritional balance between collected pollen and produced bees. These data cannot be used to assess whether the bees’ body composition follows the pollens’ content profiles, as we did not analyze honey bee samples at all test sites. In addition, not every potentially lacking nutrient, including essential amino acids, minerals and vitamins, was monitored in our field study. Nonetheless, the current study provides insight into the temporal and spatial fluctuations of pollen-derived nutrients for honey bee colonies.

The amount of protein expected in the honey bee’s body mass was lower than the amount in the collected pollen (Fig. 3A). This difference can be explained by the efficiency of pollen digestion, as a certain extent remains undigested, and thus not all of the protein in pollen is absorbed by young nurse bees (Crailsheim et al., 1992). The expected total FA content of the honey bee body mass was sometimes higher than the actual amount of total FA coming into the colonies via pollen nutrition (Figure 3B). This indicates that bees biosynthesize certain nonessential FAs, either as nurses while processing the pollen into worker jelly or during development as larvae, pupae, or young bees (Feldlaufer et al., 1985; Svoboda et al., 1982).

Unlike the storage of honey, which can reach many kilograms, bees have been reported to store only up to about 1 kg of pollen (Seeley, 1995). The colonies in the current study stored similar pollen amounts over all sites (Avni et al., 2009), albeit with a seasonal pattern paralleling the amounts of open and capped brood (Supplement 1). This suggests that in bees, pollen-intake targets and pollen storage are brood-dependent.

The proportion of linolenic acid in bee-collected pollen was generally higher than that of linoleic acid (Fig. 2). There was a distinct discrepancy in the essential FA content in Hofit (see graphical abstract). In this context, it is interesting that colonies sometimes collected relatively high amounts of particular nutrients (Fig. 3), but this was not translated into the production of more bees. However, high uptake of a specific nutrient may be a side effect of compensation for deficiency in another nutrient. The limiting nutrient in this case could be linoleic acid, as the other sites were relatively low in linoleic acid in comparison to Hofit (Fig. 2). Hence, to achieve their target level of linoleic acid, colonies may have increased pollen uptake. That linoleic acid might be a limiting factor to growth levels is supported by the production cost per bee, having the lowest CV (Table 5).

Knowing which dietary ingredient is in short supply may allow beekeepers to supplement diets with the lacking nutrient when suboptimal growth conditions are encountered (e.g., when using honey bee hives for pollination in greenhouses). Food supplementation is a common practice in animal nutritional sciences (Leeson and Summers, 2001; Lupatsch et al., 2001), and we believe that it could be adopted more extensively in apiculture as well. For example, we have shown that linolenic acid levels can be relatively low in Israel, either spatially, such as in Hofit, or temporally, in autumn (Table 2). The practice of applying omega 3 supplements to colonies might mitigate the chances of a potential nutritional deficiency for this essential fatty acid.

In summary, in support of our first hypothesis, we found that colony uptake of pollen can differ both spatially and temporally for pollen quantity and content. We also found support for altitude, as a measure of the growth season, affecting honey bee colony growth. The relatively high numbers of bees produced per year in our study in a subtropical climate compared to those reported in more temperate zones suggest that mild winter conditions can greatly increase annual honey bee production. This was further supported in our study, where colonies at higher elevations produced fewer bees than those at lower elevations. This effect may have important consequences in the face of global climate changes. We also found that the queens’ peak egg-laying rate is about double that of the mean rate calculated over the full year. This shows that total honey bee production is limited by additional factors. Our findings do not support the hypothesis of pollen quantity and nutritional quality affecting honey bee colony growth, at least not on an annual basis. For example, the site with the highest colony growth had the lowest uptake of pollen and amounts of protein and FAs (Table 3). However, it seems that relative to other reports, our colonies generally collected relatively high amounts of total protein, and were consequently able to produce a relatively large number of bees. When colonies can forage from relatively diverse landscapes, they may be able to acquire the needed levels of essential nutrients. The availability of particular nutrients may be low in particular locations or seasons, but severe nutritional deficiencies are more likely in especially depleted environments, for example in agricultural monocultures or in greenhouses.

Whether bees can then discriminate between pollens based on their nutritional composition and compensate for such deficiencies deserves further study.

## Acknowledgements

This study was funded by MERC Grant No. TA-MOU-03-M22-023, and by a grant from BBSRC, Defra, NERC, the Scottish Government and the Wellcome Trust, under the Insect Pollinators Initiative. We wish to thank the cooperating beekeepers: Eitan Zion from Yad Mordechay apiaries, Menachem Tur and Uri Surkin. We also thank Ofra Kedar for protein and fatty acid analyses, Yosi Slabetsky, Haim Kalev, Dani Barkai, Mario Ryfa, Orit Rot, Erez Tsur and Karmi L. Oxman for their technical assistance.

## Supplement 1

**Fig. S1.**
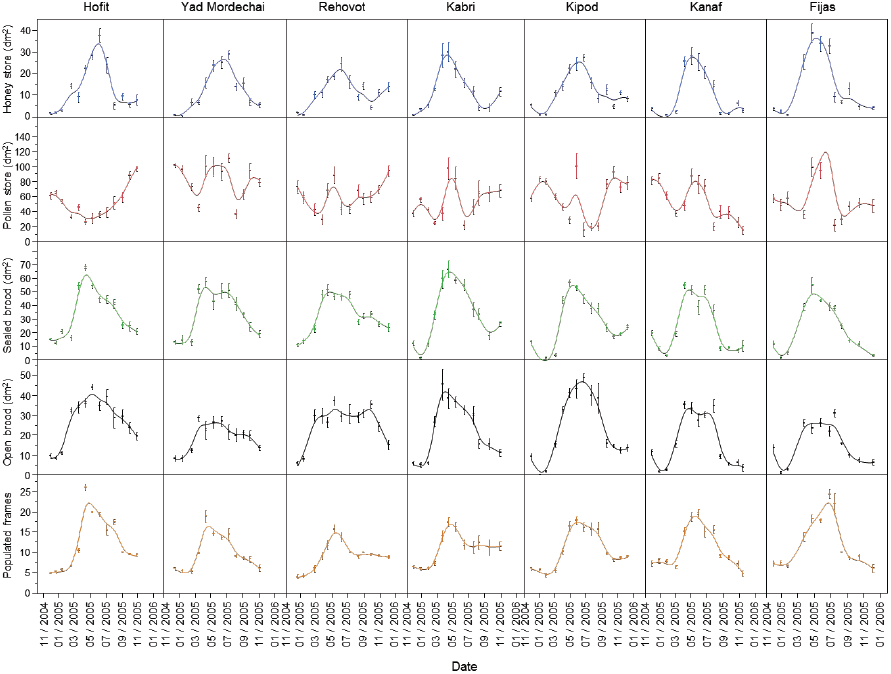
Over the course of one year, 802 hive assessments were performed: 70 hives over 7 sites with an average of 12 samplings per colony. Below are the average results of five variables for seven sites over the year, showing standard errors over the different colonies per sampled site and test date. The curves are a LOESS fit. The variables were given for total comb side coverage per colony (dm^2^): pollen stores (blue), honey stores (red), open brood (black) and closed brood (green). The bee population was assessed by the number of occupied frames per hive (yellow).

### Methodology

Standard procedures were followed, in which one person communicated observations to an assistant who recorded the data on a standard worksheet. Every comb side within a colony was closely observed. The number of worker cells filled with either stored honey or stored pollen, open brood or closed brood was assessed by estimating total comb side coverage to the nearest 0.5 dm^2^ for each of these variables. For example, when a 5 dm^2^ surface of capped brood was observed, though with an estimated 10% empty cells, the total capped brood amount was recorded as 4.5 dm^2^ coverage for that comb side. Such estimates were performed for every parameter on each comb side. In addition, the total number of frames populated by bees, and whether the colony had a notable condition such as observable disease symptoms, were noted.

## References

Alaux, C., Ducloz, F., Crauser, D., Le Conte, Y., 2010. Diet effects on honeybee immunocompetence. Biology Letters 6, 562–565.

Alfonsus, E.C., 1933. Zum Pollenverbrauch des Bienenvolkes. Arch Bienenkunde 14, 220–223.

Amdam, G.V., Page, R.P., 2005. Intergenerational transfers may have decoupled physiological and chronological age in a eusocial insect. Ageing Research Reviews 4, 398–408.

Avni, D., Dag, A., Shafir, S., 2009. Pollen sources for honeybees in Israel: Source, periods of shortage, and influence on population growth. Israel Journal of Plant Sciences 57, 263–275.

Babendreier, D., Kalberer, N., Romeis, J., Fluri, P., Bigler, F., 2004. Pollen consumption in honey bee larvae: a step forward in the risk assessment of transgenic plants. Apidologie 35, 293–300.

Bradford, M.M., 1976. Rapid and sensitive method for quantitation of microgram quantities of protein utilizing pronciple of protein-dye binding. Analytical Biochemistry 72, 248–254.

Brodschneider, R., Crailsheim, K., 2010. Nutrition and health in honey bees. Apidologie 41, 278–294.

Canavoso, L.E., Jouni, Z.E., Karnas, K.J., Pennington, J.E., Wells, M.A., 2001. Fat metabolism in insects. Annual Review of Nutrition 21, 23–46.

Cohen, A.C., 2004. Insect Diets Science and Technology. CRC Press, Boca Raton, Florida.

Crailsheim, K., Schneider, L.H.W., Hrassnigg, N., Bühlmann, G., Brosch, U., Gmeinbauer, R., Schöffmann, B., 1992. Pollen consumption and utilization in worker honeybees (*Apis mellifera carnica*): Dependence on individual age and function. Journal of Insect Physiology 38, 409–419.

de Arruda, V.A.S., Pereira, A.A.S., de Freitas, A.S., Barth, O.M., de Almeida-Muradian, L.B., 2013. Dried bee pollen: B complex vitamins, physicochemical and botanical composition. Journal of Food Composition and Analysis 29, 100–105.

Dimou, M., Thrasyvoulou, A., 2007. Seasonal variation in vegetation and pollen collected by honeybees in Thessaloniki, Greece. Grana 46, 292–299.

Douglas, A.E., 2013. Microbial Brokers of Insect-Plant Interactions Revisited. Journal of Chemical Ecology 39, 952–961.

Dussutour, A., Simpson, S.J., 2009. Communal Nutrition in Ants. Current Biology 19, 740–744.

Feldlaufer, M.F., Herbert, E.W., Svoboda, J.A., Thompson, M.J., Lusby, W.R., 1985. Makisterone-A - the major ecdysteroid from the pupa of the honey bee, Apis mellifera Insect Biochemistry 15, 597–600.

Harbo, J.R., 1986. Effect of Population-Size on Brood Production, Worker Survival and Honey Gain in Colonies of Honeybees. Journal of Apicultural Research 25, 22–29.

Haydak, M.H., 1934. Changes in total nitrogen content during the life of the imago of the worker honeybee. Journal of Agricultural Research 49, 21–28.

Haydak, M.H., Vivino, A.E., 1950. The changes in the thiamine, riboflavin, niacin and pantothenic acid content in the food of female honeybees during growth with a note on the vitamin K activity of royal jelly and beebread. Annals of the Entomological Society of America 43, 361–367.

Herbert, E.W., 1999. Honey Bee Nutrition, in: Graham, J.M. (Ed.), The Hive and the Honey Bee. Dadant & Sons, Hamilton, Illinois, pp. 197–233.

Herbert, E.W., Shimanuki, H., 1978. Chemical Composition and Nutritive-Value of Bee-Collected and Bee-Stored Pollen. Apidologie 9, 33–40.

Hilgert-Moreira, S.B., Nascher, C.A., Callegari-Jacques, S.M., Blochtein, B., 2014. Pollen resources and trophic niche breadth of Apis mellifera and Melipona obscurior (Hymenoptera, Apidae) in a subtropical climate in the Atlantic rain forest of southern Brazil. Apidologie 45, 129–141.

Hrassnigg, N., Crailsheim, K., 2005. Differences in drone and worker physiology in honeybees (*Apis mellifera*). Apidologie 36, 255–277.

Kalev, H., Dag, A., Shafir, S., 2002. Feeding pollen supplements to honey bee colonies during pollination of sweet pepper in enclosures. American Bee Journal 142, 675–679.

Khani, A., Moharramipour, S., Barzegar, M., Naderi-Manesh, H., 2007. Comparison of fatty acid composition in total lipid of diapause and non-diapause larvae of Cydia pomonella (Lepidoptera: Tortricidae). Insect Science 14, 125–131.

Kratzer, F.H., Latshaw, J.D., Leeson, S.L., Moran, E.T., Parsons, C.M., Sell, J.L., Waldroup, P.W., 1994. Nutrient requirements of Poultry, 9th ed. National Academy Press, Washington D. C.

Leeson, S., Summers, J.D., 2001. Nutrition of the Chicken, 4th ed. University Books, Guelph, Ontario, Canada.

Lupatsch, I., Kissil, G.W., Sklan, D., 2001. Optimization of feeding regimes for European sea bass Dicentrarchus labrax: a factorial approach. Aquaculture 202, 289–302.

Manning, R., 2001. Fatty acids in pollen: a review of their importance for honey bees. Bee World 82, 60–75.

Manning, R., Harvey, M., 2002. Fatty acids in honeybee-collected pollens from six endemic Western Australian eucalypts and the possible significance to the Western Australian beekeeping industry. Australian Journal of Experimental Agriculture 42, 217–223.

Mayntz, D., Raubenheimer, D., Salomon, M., Toft, S., Simpson, S.J., 2005. Nutrient-specific foraging in invertebrate predators. Science 307, 111–113.

Nurullahoglu, Z.U., Uckan, F., Sak, O., Ergin, E., 2004. Total lipid and fatty acid composition of Apanteles galleriae and its parasitized host. Annals of the Entomological Society of America 97, 1000–1006.

Odoux, J.F., Feuillet, D., Aupinel, P., Loublier, Y., Tasei, J.N., Mateescu, C., 2012. Territorial biodiversity and consequences on physico-chemical characteristics of pollen collected by honey bee colonies. Apidologie 43, 561–575.

Page, R.E., Metcalf, R.A., 1984. A population investment sex-ratio for the honey bee (*Apis mellifera* L). American Naturalist 124, 680–702.

Pirk, C.W.W., Boodhoo, C., Human, H., Nicolson, S., 2009. The importance of protein type and protein to carbohydrate ratio for survival and ovarian activation of caged honeybees (Apis mellifera scutellata). Apidologie 41, 62–72.

Raubenheimer, D., Simpson, S.J., 1999. Integrating nutrition: a geometrical approach. Entomologia Experimentalis et Applicana 91, 67–82.

Rock, G.C., King, K.W., 1967. Estimation by carcass analysis of growth requirements for amino acids in codling moth *Carpocapsa pomonella* (Lepidoptera - Olethreutidae). Annals of the Entomological Society of America 60, 1161–&.

Rosov, S.A., 1944. Food consumption by bees. Bee World 25, 94–95.

Roulston, T.H., Cane, J.H., 2000. Pollen nutritional content and digestibility for animals. Plant Systematics and Evolution 222, 187–209.

Roulston, T.H., Cane, J.H., Buchmann, S.L., 2000. What governs protein content of pollen: Pollinator preferences, pollen-pistil interactions, or phylogeny? Ecological Monographs 70, 617–643.

Schmickl, T., Crailsheim, K., 2001. Cannibalism and early capping: strategy of honeybee colonies in times of experimental pollen shortages. Journal of Comparative Physiology a-Sensory Neural and Behavioral Physiology 187, 541–547.

Schmidt, J.O., 1984. Feeding Preferences of Apis-Mellifera L (Hymenoptera, Apidae) - Individual Versus Mixed Pollen Species. Journal of the Kansas Entomological Society 57, 323–327.

Schmidt, J.O., Buchmann, S.L., 1999. Other products of the hive, in: Graham, J.M. (Ed.), The Hive and the Honey Bee, Hamilton IL, pp. 928–977.

Schmidt, J.O., Thoenes, S.C., Levin, M.D., 1987. Survival of honey bees, *Apis mellifera* (Hymenoptera, Apidae), fed various pollen sources. Annals of the Entomological Society of America 80, 176–183.

Schmidt, L.S., Schmidt, J.O., Rao, H., Wang, W.Y., Xu, L.G., 1995. Feeding Preference and Survival of Young Worker Honey-Bees (Hymenoptera, Apidae) Fed Rape, Sesame, and Sunflower Pollen. Journal of Economic Entomology 88, 1591–1595.

Seeley, T.D., 1995. The wisdom of the hive: The social physiology of honey bee colonies. Harvard University Press, Cambridge, Massachusetts.

Shafir, S., Dag, A., Haddad, N., 2009. Report: Improving honey bee colony performance by feeding pollen supplements, MERC Grant No. TA-MOU-03-M22-023. The Hebrew University of Jerusalem, Rehovot, 76100, Israel.

Simpson, S.J., Sibly, R.M., Lee, K.P., Behmer, S.T., Raubenheimer, D., 2004. Optimal foraging when regulating intake of multiple nutrients. Animal Behaviour 68, 1299–1311.

Sklan, D., Budowski, P., 1979. Cholesterol metabolism in the liver and intestine of the chick: effect of dietary cholesterol, taurocholic acid and cholestyramine. Lipids 14, 386–390.

Svoboda, J.A., Thompson, M.J., Herbert, E.W., Shortino, T.J., Szczepanikvanleeuwen, P.A., 1982. Utilization and metabolism of dietary sterols in the honey bee and the yellow-fever mosquito. Lipids 17, 220–225.

Tüylü, A.Ö., Sorkun, K., 2006. Protein analysis with kjeldahl of pollen grains collected by *Apis mellifera* L. Mellifera 6, 7–11.

Vanbergen, A.J., Baude, M., Biesmeijer, J.C., Britton, N.F., Brown, M.J.F., Brown, M., Bryden, J., Budge, G.E., Bull, J.C., Carvel, C., Challinor, A.J., Connolly, C.N., Evans, D.J., Feil, E.J., Garratt, M.P., Greco, M.K., Heard, M.S., Jansen, V.A.A., Keeling, M.J., Kunis, W.E., Marris, G.C., Memmott, J., Murray, J.T., Nicolson, S.W., Osborne, J.L., Paxton, R.J., Pirk, C.W.W., Polce, C., Potts, S.G., Priest, N.K., Raine, N.E., Roberts, S., Ryabov, E.V., Shafir, S., Shirley, M.D.F., Simpson, S.J., Stevenson, P.C., Stone, G.N., Termansen, M., Wright, G.A., Initiative, I.P., 2013. Threats to an ecosystem service: pressures on pollinators. Frontiers in Ecology and the Environment 11, 251–259.

Vanderplanck, M., Leroy, B., Wathelet, B., Wattiez, R., Michez, D., 2014. Standardized protocol to evaluate pollen polypeptides as bee food source. Apidologie 45, 192–204.

Wang, Y.M., Lin, D.S., Bolewicz, L., Connor, W.E., 2006. The predominance of polyunsaturated fatty acids in the butterfly Morpho peleides before and after metamorphosis. Journal of Lipid Research 47, 530–536.

Wille, H., Wille, M., Kilchenmann, V., Imdorf, A., Buhlmann, G., 1985. Pollen Gathering and Population-Dynamics of 3 Liebefeld Bee Colonies of Apis-Mellifera during 2 Consecutive Years. Revue Suisse De Zoologie 92, 897–914.

Winston, M.L., 1987. The Biology of the Honey Bee. Harvard University Press ISBN 0-674-07408-4.

Wright, W., 2008. How many eggs can a queen lay? Bee Culture, November 2008, pp. 59–61.

Yang, K., Wu, D., Ye, X.Q., Liu, D.H., Chen, J.C., Sun, P.L., 2013. Characterization of Chemical Composition of Bee Pollen in China. Journal of Agricultural and Food Chemistry 61, 708–718.

